# Hidden state models improve the adequacy of state-dependent diversification approaches using empirical trees, including biogeographical models

**DOI:** 10.1101/302729

**Authors:** Daniel S. Caetano, Brian C. O’ Meara, Jeremy M. Beaulieu

## Abstract

The state-dependent speciation and extinction models (SSE) have recently been criticized due to their high rates of “false positive” results and many researchers have advocated avoiding SSE models in favor of other “non-parametric” or “semi-parametric” approaches. The hidden Markov modeling (HMM) approach provides a partial solution to the issues of model adequacy detected with SSE models. The inclusion of “hidden states” can account for rate heterogeneity observed in empirical phylogenies and allows detection of true signals of state-dependent diversification or diversification shifts independent of the trait of interest. However, the adoption of HMM into other classes of SSE models has been hampered by the interpretational challenges of what exactly a “hidden state” represents, which we clarify herein. We show that HMM models in combination with a model-averaging approach naturally account for hidden traits when examining the meaningful impact of a suspected “driver” of diversification. We also extend the HMM to the geographic state-dependent speciation and extinction (GeoSSE) model. We test the efficacy of our “GeoHiSSE” extension with both simulations and an empirical data set. On the whole, we show that hidden states are a general framework that can generally distinguish heterogeneous effects of diversification attributed to a focal character.

## Introduction

Determining the impact that trait evolution has on patterns of lineage diversification is a fundamental and core question in evolutionary biology. The state speciation and extinction (SSE; Maddison et al. 2007) framework was developed specifically for these purposes, as it provides a means of correlating the presence or absence of a particular character state on diversification rates. Since the initial model was published, which modeled the evolution of a single binary character (i.e., BiSSE; Maddison et al. 2007, FitzJohn et al. 2009), the SSE framework has been expanded to deal with multiple state/traits (MuSSE: FitzJohn 2012), continuous traits (QuaSSE: FitzJohn, 2010), to test whether change happens at speciation events or along branches (ClaSSE: Goldberg and Igic 2012; BiSSE-ness: Magnuson-Ford and Otto 2012), which also includes a nested subset of models that examines geographic range evolution (GeoSSE: Goldberg et al. 2011), and to account for “hidden” states that may influence diversification, on their own or in combination and possible interaction with observed states (HiSSE; Beaulieu and O’Meara 2016).

The initial wave of interest and use of SSE models is quickly being replaced with widespread skepticism about their use (see O’Meara and Beaulieu 2016). One major reason is based on the simulation study of Rabosky and Goldberg (2015). Their analyses showed that if a tree evolved under a heterogeneous branching process, completely independent from the evolution of the focal character, SSE models will almost always return high support for a model of trait-dependent diversification. From an interpretational standpoint, this is certainly troubling. However, this also stems from the misconception that any type of SSE model is a typical model of trait evolution like in, say Pagel (1994) or Butler and King (2004), where the likelihood function maximizes the probability of the observed trait information at the tips, given the model *and* the tree. In these models, the phylogenetic tree certainly affects the likelihood of observing the traits, but that is the only role it plays. Other models based on the birth-death process for understanding tree growth and shape (e.g., Nee et al. 1994, Rabosky and Lovette 2008, Morlon et al. 2011) only calculate the likelihood of the tree itself, ignoring any and all traits. An SSE model is essentially a combination of these: it computes the joint probability of the observed states at the tips *and* the observed tree, given the model. This is an important distinction because if a tree violates a single regime birth-death model due to trait-dependent diversification, mass extinction events, maximum carrying capacity, or other factors, then even if the tip data are perfectly consistent with a simple trait evolution model, the tip data *plus* the tree are not. In such cases the SSE model is very wrong in assigning rate differences to a neutral trait, but it is also wrong in saying that the tree evolved under unchanging speciation and extinction rates. This leaves practitioners in quite a bind because the “right” model is not something that could be tested in the SSE framework.

These results have created continued concerns in the community with respect to SSE models. There are reasons to be concerned, but, in our view, there is a deeper issue with the misinterpretation of hypothesis testing. First, with any comparative model, including SSE, rejecting the “null” model does not imply that the alternative model is the true model. It simply means that the alternative model fits *less* badly. Misunderstanding of null hypothesis testing, and its dubious utility, has been a prominent issue for decades in other fields (i.e., Berkson 1938, Kirk 1996). Second, in biological examples, including many of those used for testing Type I error (e.g., Rabosky and Goldberg 2015), the apparent issues with SSE models isn’t a matter of high Type I error rate at all, it is simply comparing a trivial “null” model (i.e., equal rates diversification) against a model of trait-dependent diversification. Again, given the rich complexity of processes affecting diversification (e.g., mass extinctions, local extinctions, competition, and biogeographic changes) and trait evolution (e.g., varying population size, selection pressure, and available variation), a comparison of “one rate to rule them all” versus “something a bit more complex” will usually return the latter as a better descriptor of the data. A fairer comparison involves a “null” model that contains the same degree of complexity in terms of the number of different classes of diversification rates that are also independent of the evolution of the focal character, to allow for comparisons among trait-dependent models of interest (Figure 1; also Beaulieu and O’Meara 2016). The development of the hidden state SSE model (HiSSE; Beaulieu and O’Meara 2016) provides a means of including unobserved “hidden” characters into the model that can account for the differences in diversification.

**Figure 1.**
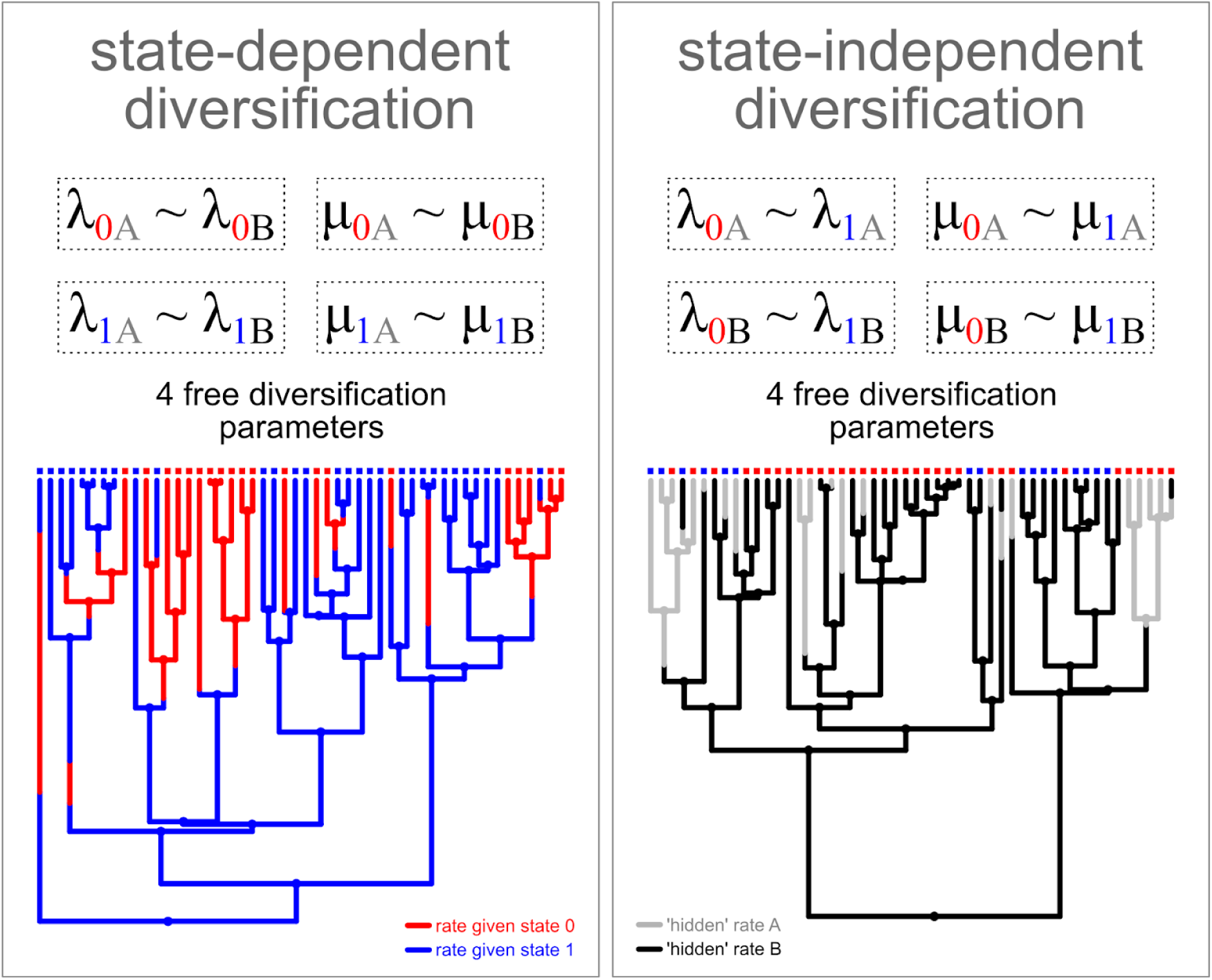
Hidden state models provide a framework to solve the issue with false relationships between shifts in diversification rates and traits. Parameters linked by ‘∼’ are constrained to share the same value. Left-side panel shows a case of state-dependent evolution. Here, shifts in rates of diversification in the tree are perfectly predicted by the transition between trait states 0 (red) and 1 (blue). Right-side panel shows state-independent shifts in diversification rates with respect to the focal trait (gray vs. black branches). Both models share the same number of free diversification parameters, but the variation among hidden states is partitioned differently. The state-independent model (right panel) allows for two diversification rate categories unrelated to the observed traits. An overly simplistic homogeneous rate model would be inadequate for either of these trees.

Aside from issues related to model rejection, and what is the appropriate “null” comparison, there are also broader issues related to any interpretation of trait-dependent diversification. For instance, a typological flaw in these types of analyses is the implicit assumption that a single trait is the primary driver of diversification, thereby ignoring alternate sources of rate variation. In reality, nearly all traits exert at least *some* effect on speciation and/or extinction rates. Even something as modest as a different base at a third codon position that leads to the same amino acid can have a tiny fitness difference (Yang and Nielsen 2008), which in theory could make a species infinitesimally closer to extinction. Moreover, even among traits we think have a greater effect, it is unlikely that *only* this one examined trait accounts for the increased diversification rates. In other words, if we are studying, say, growth habit, it is very unlikely that traits like floral symmetry, pollination syndrome, biogeography, fruit dispersal, etc. all have exactly zero effect on diversification. It should, therefore, always be difficult to ever confidently view any one character state as the true underlying cause of changes in diversification rates. The inclusion of “hidden” traits, again, provides a means of testing intuitions about a particular character, while also “soaking” up variation in diversification rates that is also driven by some unknown or unmeasured factor (which may interact with the particular character, as well).

The use of “hidden” states, and the hidden Markov modelling approach (HMM) in general, addresses multiple issues at once and are fairly simple to implement in an SSE framework. They are, however, challenging from a interpretational standpoint. For example, what does a “hidden” state mean? How does one weigh evidence for or against trait-dependent and trait-independent diversification in the presence of “hidden” rates? The purpose of this study is twofold. First, given the confusion over whether SSE models remain a viable means of assessing state-dependent diversification (e.g., Rabosky and Goldberg 2017), we further clarify the concept of “hidden” states as well as the misconceptions of “Type I errors”. Second, we demonstrate the role of hidden state models as a general framework by expanding the original geographic state speciation and extinction model (GeoSSE; Goldberg et al. 2011). We define a set of biologically meaningful models of area-independent diversification (AIDiv) to be included in studies of area-dependent diversification (ADDiv) that can be used in combination with the original GeoSSE formulation. Such models are especially useful given the recent clarification on the undesirable impact of cladogenetic events on the performance of dispersal-extinction-cladogenesis models (DEC; Ree and Smith 2008 and DEC+J; Matzke 2014) as well as the renewed advocacy of using SSE models when examining patterns of geographic range evolution (Ree and Sanmartín 2018).

### The value of incorporating “hidden” states into SSE models

As mentioned above, SSE models are routinely criticized on the grounds that they almost always show increased levels of “Type I error” (Rabosky and Goldberg 2015). That is, when fitted to a tree evolved under a heterogeneous branching process independent from the evolution of the focal character, SSE models will almost always return high support for a trait-dependent diversification model over a trivial “null” model that assumes equal diversification rates. However, as pointed out by Beaulieu and O’Meara (2016), this particular issue does not represent a case of Type I error, but, rather, a simple problem of rejecting model *X*, for model *Y*, when model *Z* is true. Furthermore, rejecting model *X* for model *Y*, does not imply that model *Y* is the true model. It simply means that model *Y* is a better *approximation* to model *Z*, than model *X.* This will be generally be true if model *X* is overly simplistic (i.e., diversification rates are equal) with respect to the complexity in either model *Y* and model *Z* (i.e., diversification rates vary).

The story of the boy that cried wolf is a popular mnemonic for understanding what we mean when we refer to the difference between true Type I and Type II error, which can be extended to include comparisons between complex and overly simplistic models. When the boy first cried wolf, but there was no wolf, he was making a Type I error ‐‐ that is, falsely rejecting the null of a wolf-free meadow. When the townspeople later ignored him when there was actually a wolf, they were making a Type II error. If the sheep were instead perishing in a snowstorm, and the only options for the boy are to yell “no wolf!” or “wolf!” it is not clear what the best behavior is ‐‐ “no wolf” implies no change in sheep mortality rates from when they happily gambol in a sunny meadow, even though they have begun to perish, while “wolf” communicates the mortality increase even though it is the wrong mechanism. It is the same here when looking at a tree coming from an unknown, but complex empirical branching process and trying to compare a constant rate model (“no wolf”) against a trait-dependent (“wolf”), age-dependent (“bear”), or density-dependent model (“snowstorm”).

Beaulieu and O’Meara (2016) proposed a set of character-independent diversification (CID) models that are parameterized so that the evolution of each observable character state is independent of the diversification process without forcing the diversification process to be constant across the entire tree. Importantly, hidden state models are part of a more general framework that should be applied to any SSE model, regardless of *a priori* interest in unobserved factors driving diversification. For instance, the likelihood of a standard BiSSE model is *identical* to a HiSSE model where the observed states each have their own unique diversification rates, with the underlying “hidden” states constrained to have the same parameter values. This is best illustrated in Figure 1. Both phylogenetic trees show variation in diversification rates, so the question is whether such rate shifts can be predicted by the observed traits or not. The state-dependent and independent models with respect to a focal trait both have two hidden states (*A* and *B*) and four free diversification parameters. The difference between the models in Figure 1 resides solely in the way the variation among the hidden states is partitioned. If we set the hidden rates to vary within each observed state, such that all hidden states of λ_0_ and μ_0_ are independent of λ_1_ and μ_1_ (Figure 1, left panel), we produce a state-dependent BiSSE model. Alternatively, if the observed states share the same diversification rate, and rate variation is partitioned among hidden rate classes (Figure 1, right panel), we produce a state-independent model, which is independent of the focal state, but not the hidden state. More complex state-dependent models can be created by letting multiple hidden rate categories be estimated within the observed state (see examples in Beaulieu and O’Meara 2016). Of course, in the case of BiSSE (as well as any other of the original SSE implementations), there is little sense in referring to the single observed rate category as “hidden”, since the hidden states are constrained to have the same parameter values. Nevertheless, the usefulness of the state-independent models (CID) is that they partition the rate variation among hidden rate categories and not among observed states. For a fairer comparison to a state-dependent diversification model, any alternative CID model must be devised to fit the same number of diversification rate categories to the phylogeny (i.e., same number of free speciation and extinction rates, such as shown in Figure 1).

A natural corollary, then, is that the usefulness of the hidden state modelling approach depends entirely on the careful match of free diversification parameters between the state-dependent and state-independent diversification models. These models will also return “false positives” when the proper counter balance to a trait-dependent model is not included in the set of models evaluated. This was recently demonstrated by Rabosky and Goldberg (2017). Under a range of very difficult scenarios, their non-parametric approach differentiated between scenarios of trait-dependent and trait-independent diversification much better than a parametric, process-based hidden state SSE model. Specifically, they found that while the inclusion of a character-independent model with two diversification rate categories (i.e., CID-2) reduced the overall “false positive” rate of BiSSE, the use of BiSSE + CID-2 + HiSSE resulted in nine of 34 trait-independent diversification scenarios having “false-positive” rates in excess of 25%. The results of Rabosky and Goldberg (2017) show that appropriate null models help, but they are not a panacea. However, we hasten to point out that this result is partly due to not including sufficiently complex null models that are able to capture enough variation in diversification rates across the phylogenies. In fact, the most parameter-rich model tested by Rabosky and Goldberg (2017) assumed trait-dependent diversification. A proper set of CID models for hidden state SSE methods will, necessarily, have the same number of free diversification parameters than the state-dependent models (the CID-4 in this case). When proper null models are included, the “false positive” rates dropped in all scenarios (see Figure 2). Moreover, the improvement in performance was dramatic in the same nine scenarios that previously showed high “false-positive” rates (see Figure 6 in Rabosky and Goldberg 2017). Thus, not just any null model will suffice, but once appropriately complex ones are included, Type I properties approach desired values.

**Figure 2.**
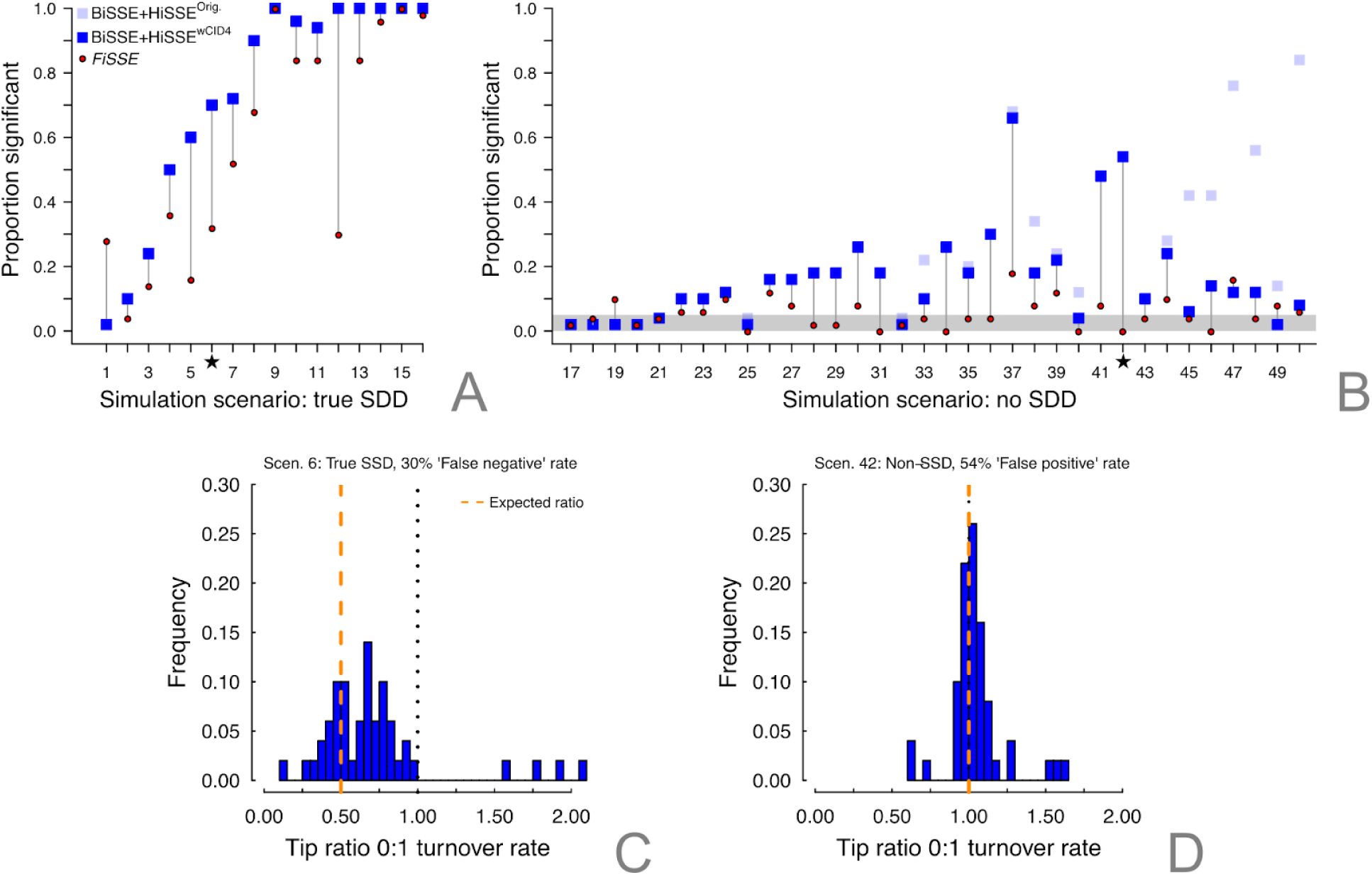
Reanalysis of the Rabosky and Goldberg (2017) when the 4-state character independent model, CID-4, is included in the model set (dark blue boxes). When compared against the fit of BiSSE, CID-2, and HiSSE, the (A) power to detect the trait-dependent diversification remains unchanged. For the trait-independent scenarios (B), there is almost always a reduction in the “false positive” rate (as indicated by the difference in the position of the light blue and dark blue boxes), and in many cases the reduction is substantial. When we model-averaged parameter estimates of turnover from two scenarios, a true SSD scenario (C), where the HiSSE model set showed high “false negative” rates (i.e., failed to reject a trait-independent scenario), and a non-SSD scenario (D) which exhibited a 54% false-positive rate. In the case of the non-SSD scenario, it clearly shows that despite the poor performance of from a model rejection perspective, examining the the model parameters indicates that, on average, there are no differences in diversification rates among observed states. The dashed orange line represents the expected ratio to be compared against a ratio of difference in diversification rates between state 0 and 1 denoted by the dotted black line.

### What (if anything) is a hidden character?

Common difficulties that come with hidden Markov models applied to comparative analyses are “what exactly does the hidden state represent?”, which leads to the most pressing question of “how should I interpret results when I find evidence for one or more hidden states influencing diversification?” Before moving forward, it is important to note a continuum across models of traits and diversification, such as the SSE models, and those that fit diversification rates to trees but ignore character information altogether (LASER, Rabosky 2006; MEDUSA, Alfaro et al. 2009; TreePar, Stadler 2011; BAMM, Rabosky 2014; among others). While all these models are often treated as unrelated frameworks, they are really two ends of a continuum. On one end lie models such as MEDUSA and BAMM that make no explicit hypotheses about how traits impact diversification, but implicitly assume the inferred shifts must be tied to something about the organism or its environment. In fact, these models can be considered “hidden state only” models, in that shifts in diversification could be related to a single unmeasured character or, more realistically, to an evolutionary coordination among a suite of traits and trait-environment interactions. In other words, it is not controversial to assert that most characters have at least some influence on diversification, however trivial, and that even when not explicitly identifying a character focus, we are doing so implicitly. This is precisely why trait-based interpretations come as part of post-hoc interpretations that assert the putative causality of shifts in diversification inferred with MEDUSA or BAMM.

On other end of the continuum, we may have a hypothesis about particular character states and their impact on diversification, which we test with any one of the many available -SSE models. However, such questions increasingly now come with the added burden of mitigating factors that may erroneously produce meaningful differences in diversification among character states (Maddison and FitzJohn 2015, Beaulieu and O’Meara 2016), which ultimately leads us towards a middle ground of blending tree-only and strict -SSE type models. For instance, as mentioned above, we must account for the possibility that diversification rates may actually vary independently of our special character of interest. Even when diversification rates are tied to a particular focal character state, we must also account for complicated correlations between our trait of interest with one or more unmeasured characters that can vary among clades (Beaulieu and O’Meara 2016). We should also account for additional processes, such as whether speciation events (cladogenesis) exert an effect on the character even if it may seem inconsistent with our initial hypothesis of how a particular state evolves. The important point here is that without accounting for any or all of these factors, and by explicitly staying on the strict end of the -SSE spectrum, we run the risk of overstating the meaningfulness of the diversification differences among character states.

So, what, if anything, is the purpose of a hidden state model? Simply put, it is a means by which we can account for the hidden majority of traits while examining the meaningful impact of a suspected “driver” of diversification. The nature of the empirical question of whether character state *X* has a meaningfully higher rate than character state *Y* does not, and should not, change when including hidden characters. However, we emphasize that this type of question serves no purpose if character states *X* and *Y* alone are not adequate predictors for diversification rates. In this case, neither the “null” model, where diversification rate for *X* and *Y* are the same, nor the alternative model of trait-dependent diversification will be adequate. If we ignore within trait variation in diversification rates, it is really not surprising that erroneous conclusions are made. For these reasons, we advocate moving beyond just rejecting trivial null models, by making comparisons among a variety of models, looking at the weight for each, and making biological conclusions based on some summary of these models and their parameter estimates.

### The importance of model-averaging

We adopt an approach that integrates the estimate from every model in the set in proportion to how much the model is able to explain the variance in the observed data (i.e., model-averaging by Akaike weight, *w*_i_). Although the Akaike weights can also be used to rank and select the “best” model(s), the fundamental difference between approaches is that model-averaging largely alleviates the subjectivity of choosing thresholds to rank models, and permits the estimation of parameters taking into account the uncertainty in model fit. For instance, it is not unusual for a set of models, ranked according to their AIC values, to suggest that three or four models fit about “equally well”. The model choice framework provides no easy solution for such instances and, importantly, the conclusions about the biological questions at hand end up hampered by potentially conflicting interpretations from the models. Parameter estimates averaged across these models, on the other hand, will result in a unique scenario which incorporates the uncertainty associated with the fit of multiple models. For example, in the case of the data sets simulated by Rabosky and Goldberg (2017), the scenario that exhibited one of the worst “false positive” rates, even after properly accounting for the CID-4 in the model set, had a model-averaged tip ratio between observed states, *0* and *1*, distributed around 1 (see Figure 2). This indicates that, on average, there were no meaningful diversification rate differences among the observed states. More importantly, this is a clear example of a case in which “false-positive” results are not as dire as they seem, because parameter estimates for the model show no strong effect of state-dependent diversification (see similar discussion in Cooper et al. 2016b).

The procedure to perform parameter estimates starts by computing the marginal probability of each ancestral state at an internal node using a standard marginal ancestral state reconstruction algorithm (Yang et al. 1995, Schluter et al. 1997), though using the SSE model rather than just a model for the trait. The marginal probability of state *i* for a focal node is the overall likelihood of the tree and data when the state of the focal node is fixed in state *i*. Note that the likeliest tip states can also be estimated, because while we observe state *1*, the underlying hidden state could either be *1A* or *1B.* For any given node or tip we traverse the entire tree as many times as there are states in the model. Second, the weighted average of the likeliest state and rate combination for every node and tip is calculated under each model, with the marginal probability as the weights. Finally, the rates and states for all nodes and tips are averaged across all models using the Akaike weights. These model-averaged rates, particularly among extant species, can then be examined to determine the tendency of the diversification rates to vary among the observed character states. This procedure is implemented in the *hisse* package (Beaulieu and O’Meara 2016).

One important caveat about model-averaging is making sure the models that are being averaged provide reasonable parameter estimates. Including a model with a weight of 0.00001, but a parameter estimate millions of times higher than other models’ estimates for that parameter, will have a substantial effect on the model-average. Even with this approach, examining results carefully, and communicating any issues and decisions to cull particular models transparently, remains important.

### Similarities between BAMM and model-averaging using AIC*w*

Model-averaging approaches have been widely used in phylogenetics (e.g., Posada 2008, Eastman et al. 2011, Rabosky 2014), but most applications have focused on Bayesian methods. For example, BAMM (Rabosky 2014) applies a reversible-jump Markov-chain Monte Carlo (rjMCMC) sampler (Green 1995). The “reversible-jump” part means that the MCMC will visit multiple models by adding and subtracting parameters using birth and death proposal steps. This is a Bayesian method, thus a prior is used to control the weight given to different numbers of rate shifts (Rabosky 2014). The rjMCMC will adjust the complexity of the model in function of the likelihood weighted by the prior (i.e., the posterior). This approach returns a posterior distribution of shifts in net diversification and locations in the phylogenetic tree.

The SSE trait-independent models using hidden states (CID - Beaulieu and O’Meara 2016) have a similar purpose to the trait-agnostic analyses of rates of diversification performed with BAMM. In both cases we want to compute the likelihood of the tree including rate heterogeneity in the absence of any (explicit) predictor. As we discussed before, the important attribute of the CID models is that these can accommodate rate heterogeneity using hidden states, in contrast to simple null models, such as in BiSSE. Since the true number of rate shifts in empirical trees is unknown, we need to fit multiple trait-independent models with varying number of hidden rate categories. While BAMM has a built-in machinery to grow and shrink the number of rate regimes in the models as part of the rjMCMC, HiSSE (as well as GeoHiSSE, which we describe below) requires the user to provide a set of models with an adequate number of rate categories (i.e., hidden states). However, it is possible to dredge across a set of possible models with a simple script and summarize results using model averaging (see *Supplementary Material*). More importantly than approximating the number of rate shifts in tree, the number of free diversification rate categories of the trait-independent models always need to match the rate categories present in the trait-dependent alternative models (as shown in Figure 1).

The similarity between the application of model-averaging in a likelihood framework and Bayesian model-averaging is that, in both cases, the result is a collection of models weighted by some quantity proportional to a measure of fit. In the case of model averaging across a set of maximum likelihood estimates, the quantity used is the Akaike weight which are considered the weight of evidence in favor of a given model relative to all other models in the set (Burnham and Anderson 2002). In a Bayesian framework, the quantity is the frequency of a given model in the posterior distribution which is proportional to the posterior probability of the model given the observed data (Green 1995). Both the rjMCMC utilized by BAMM and model averaging approach described here have the same ultimate goal and allow us to investigate similar questions. Given the recent popularity of rjMCMC approaches in phylogenetics, it seems natural that tests of state-dependent diversification using likelihood should focus on parameter estimates while incorporating the contribution of each model to explain the observed data and avoid the pitfalls of model testing.

### Linking “hidden” states to geographic range evolution and diversification

The dispersal-extinction-cladogenesis models (DEC; Ree and Smith 2008) as well as DEC+J (“jump dispersal”; Matze 2014) are popular frameworks for studying geographic range evolution using a phylogenetic-based approach. They are different from SSE models in that they are standard trait evolution models ‐‐ that is, the likelihood *only* reflects the evolution of ranges and the tree is considered fixed. Recently, Ree and Sanmartín (2018) raised concerns with the limitations of DEC and DEC+J models. Since the probability of the tree is not part of the DEC likelihood, cladogenetic events are independent of time, which produces odd behaviors in the parameter estimates, especially when jump dispersal events are allowed (Ree and Sanmartín 2018). A straightforward way to alleviate this issue, as mentioned by Ree and Sanmartín (2018), is to simply apply geographic state speciation and extinction models (GeoSSE; Goldberg et al. 2011) to study range evolution. Instead of optimizing ancestral areas conditioned on the nodes of a fixed tree, the GeoSSE model incorporates the tree into the likelihood and has a parameter for the rate of cladogenetic speciation.

Adopting an SSE-based framework to understand geographic range evolution is naturally burdened with respect to comparing complex models to “trivial” nulls and accounting for “hidden” variation among observed states. In our view, the concept of hidden variation seems the most relevant when investigating geographic range evolution. Due to the need for reducing geographical variation into coarsely defined discrete areas, parameter estimates could be strongly impacted by heterogeneous features across the landscape not captured by this categorization. A good example of the potential for hidden variation in range evolution comes from studies of diversity dynamics between tropical and temperate regions, which are often defined simply by latitude (e.g., tropical for | latitude | < 23.5 degrees; Rolland et al. 2014). Such categorization necessarily overlooks the heterogeneity present in the tropics: some high elevation areas freeze, others do not; some areas are deserts and others are lush forests, etc. However, the ability for the data to “speak” to the rate variation within a given geographic area is not currently allowed within the existing GeoSSE framework. Below, we briefly demonstrate how the hidden Markov modelling (HMM) approach can easily be used for geographical state speciation and extinction models. However, this is just one example of the utility of HMM approaches in diversification models; HMM approaches can, and we argue should, be added as options to other diversification approaches.

*Geographic hidden state speciation and extinction model* ‐‐ The general form of the original GeoSSE model determines the diversification dynamics within, and transitions between, two discrete regions *0* and *1.* Under this model (see Figure 3), a species observed at the present (*t*= 0) can be “endemic” to either area *0* or *1*, or has persistent populations in both *0* and *1*, referred to hereafter as the *01* range (i.e., “widespread”). Similarly to the DEC model, range evolution occurs in two distinct modes. First, ranges can expand or contract along the branches of the phylogeny through anagenetic change. Range expansions are based on state transitions from *0* to *01* and *1* to *01*, which are parameterized in the model as the per-lineage “dispersal” rates, *d_0_* and *d_1_* respectively. Range contractions, on the other hand, describe the reverse process of transitions from *01* to *1* and *01* to *0*, which are the per-lineage rates of range contraction, *x_0_* and *X_1_*, respectively (also referred as “extirpation” rates). The second mode of range evolution occurs as a product of the speciation process (i.e., cladogenesis), particularly with respect to speciation events breaking up widespread ranges into various combinations of descendant areas. The area-specific rates of “within-region” speciation are parameterized as *s_0_* and *s_1_* whereas the “between-region” rate of speciation is denoted by *s_01_.* We will refrain from describing the mathematical formulation of this particular model, as these are described in detail elsewhere (Goldberg et al. 2011, Goldberg and Igic 2012). We do note, however, that our notation for the observed areas differs from Goldberg et al. (2011) in order to be consistent with previous work on incorporating hidden states into -SSE models (see Beaulieu and O’Meara 2016).

**Figure 3.**
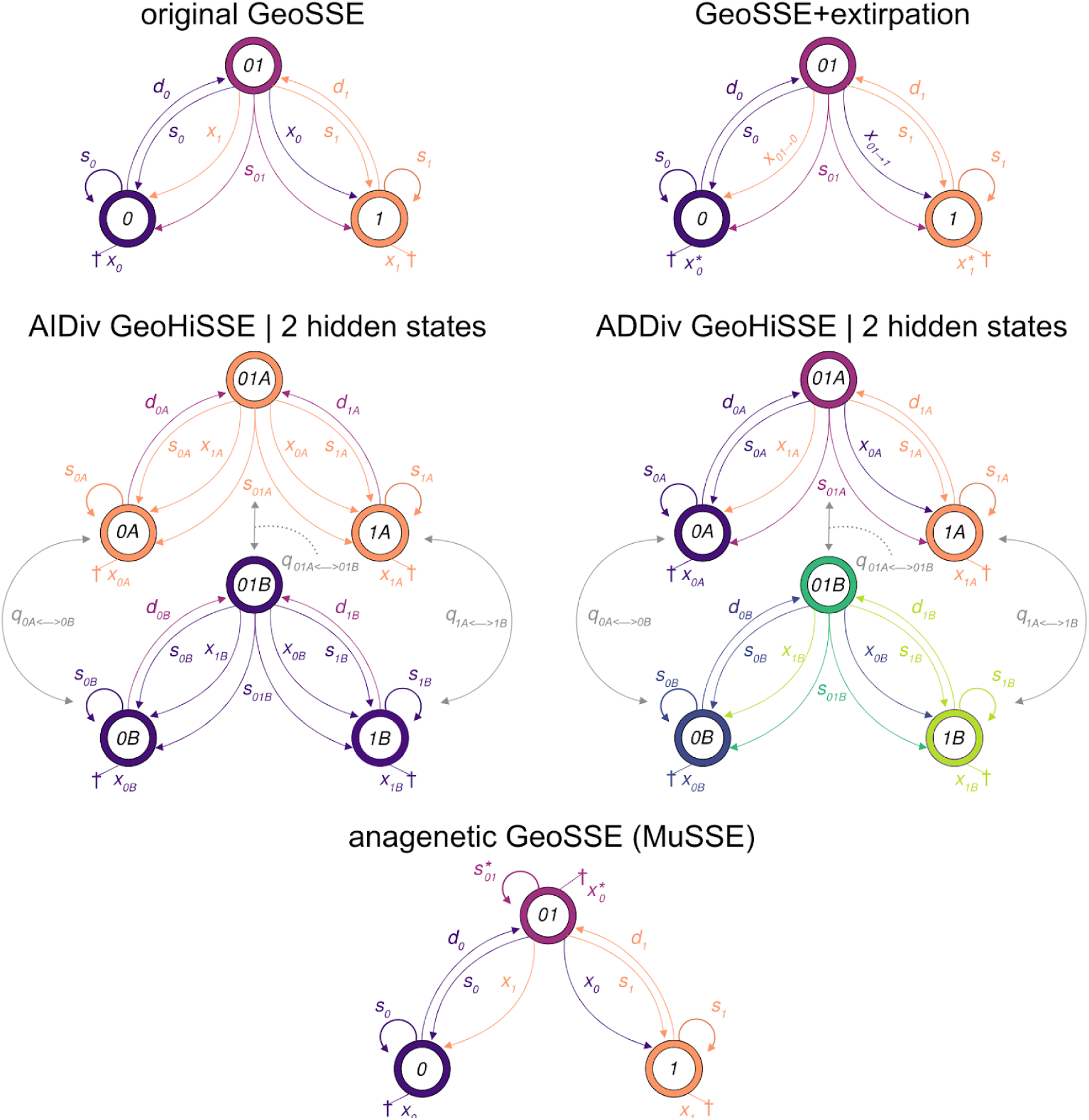
The states and allowed transitions in the original GeoSSE model (Goldberg et al., 2011) and the model extensions described herein. The colors denote parameters that are associated with each character combination modelled as having their own unique set of diversification rate parameters. The GeoSSE+extirpation model separates rates of range reduction (e.g., x_01→1_) from the extinction of endemic lineages (e.g., x_0_), but otherwise contains three unique sets of diversification parameters as in the original GeoSSE model. The area independent diversification (AIDiv) GeoHiSSE (denoted by two sets of diversification parameters shown in orange and purple) and the area dependent diversification (ADDiv) GeoHiSSE (denoted by six sets of diversification parameters shown in various colors) models can have 2 or more hidden states. Finally, the anagenetic GeoSSE model is a special case of the MuSSE model parameterized as to emulate the transitions allowed by the original GeoSSE model. Note that GeoSSE+extirpation as well as the anagenetic models can also support hidden states.

In order to apply the hidden Markov modelling (HMM) to GeoSSE for the simplest case of two hidden states, we replicate the original model across hidden states *A* and *B.* We re-parameterize the model to include six distinct speciation rates, *s_0A_*, *s_1A_*, *s_0lA_*, *s_0B_*, *s_1B_*, and *s_01B_*, and four distinct extinction rates, *x_0A_*, *x_1A_*, *x_0B_*, and *x_1B_*, allowing for two distinct net diversification rates within each range (Figure 3). Likewise, dispersal rates from area *0* (or *1*) into *01* also show separate rates for each hidden state, parametrized as *d_0A_* and *d_0B_* for area *0* or *d_1A_* and *d_1B_* for area *1.* Shifts between the hidden states within a geographic range are modelled with the transition rate *q* following the same approach described by Beaulieu and O’Meara (2016), as does the inference of changes in the hidden state within each range. Herein we focus our simulation tests and empirical analyses on a spatial structure that contains only two geographical areas, however, just like the DEC model, our GeoHiSSE model can contain an arbitrarily large state space.

The GeoSSE model set must also be expanded to include a more complex and less trivial set of “null” models to compare against those that assume some form of area-dependent diversification (referred to here as ADDiv). Recently, Huang et al. (2017) extended the CID approach to obtain an area-independent model (referred to hereafter as, AIDiv) for use in model comparisons against GeoSSE. The AIDiv model of Huang et al. (2017) replicates the GeoSSE model across two hidden states, *A* and *B*, similar to our GeoHiSSE model described above. However, here the diversification rates are constrained to be equal in each of the hidden states such that *s_0A_* = *s_1A_* =*s_01A_*, *x_0A_* = *x_1A_*, *s_0B_* = *s_1B_* = *s_01B_*, and *x_0B_* = *x_1B_* (Figure 3). There is also a global transition rate (e.g., *d*_*0A*->0B_) which accounts for transitions among the different hidden states within each range and disallows dual transitions between areas and hidden states (i.e., *d*_*0A*->1B_=*0*). We expand the AIDiv model to allow geographic ranges to be associated with as many as five different hidden states (i.e., *h***∈***A*, *B*, *C*, *D*, *E*). The AIDiv model that contains two hidden states is equivalent to the model of Huang et al (2017). It should be noted that this model contains only two diversification rate categories, which makes the AIDiv model slightly less complex than the original GeoSSE model (which has three). In any event, the purpose of these models is to prevent spurious assignments of diversification rate differences between observed areas in cases where diversification is affected by other traits. Finally, an AIDiv model with five hidden states contains as many as 10 free diversification parameters and, importantly, equals the complexity in our GeoHiSSE model.

*Further model expansions* ‐‐ We also relaxed and tested the behavior of two important model constraints within the GeoSSE framework. First, we wondered whether constraining range contraction and lineage extinction to be the same could be too restrictive, particularly when the ranges under consideration represent large geographical areas. Under the original GeoSSE model there are two parameters, *d*_*01*->*0*_ and *d*_*01*->*1*_ which denote local range contraction that are linked to extinction rates of the endemics, *x_0_* and *x_1_*, (Figure 3). However, consider a scenario where lineages have originated in a temperate region and possess a suite of traits that reduce extinction rates in this area. Movements into the tropical regions require not just getting there, but also being able to compete within this new environment. Recent attempts to disperse into the tropical zone by those lineages on the boundary separating the two areas can persist there for a time, but might eventually go locally extinct in the tropical portion. The constraint of the rate of range contraction always equalling the extinction rate of endemics prevents the detection of such dynamics. Furthermore, this will necessarily increase estimates of per-lineage extinction rates for the tropical region as a whole because of the link between range contraction and lineage extinction present in the original GeoSSE model (Goldberg et al. 2011). Here we extended the model by separating the rate of range contraction from the process of lineage extinction (see more details in Figure 3 and *Supplementary material*). More broadly, we refer to this class of models as “GeoHiSSE+extirpation” or “GeoSSE+extirpation” to represent models with and without hidden states, respectively. Removing this constraint effectively increases the number of state “transition rates” in the model. Teasing apart the effect of range reduction and extinction of endemics will likely require phylogenetic trees of large size. For instance, our simulations show parameter estimates for the hiGeoSSE+extirpation model are adequate with trees of 500 species (see *Supplementary material*).

Second, we included a complementary set of models (both AIDiv and ADDiv) that removed the cladogenetic effect from the model entirely. These models assume that lineage speciation has no direct impact on range evolution, such that all changes occur along the branches (i.e., anagenetic change). This requires the addition of a per-lineage rate of extinction and speciation for lineages in the widespread range (*x_01_* and *s_01_*, respectively) as well as range contraction being distinct from the extinction of endemics. In the absence of a hidden state, this is effectively a three-state MuSSE model (FitzJohn 2012) with transition matrix constrained such that a shift between ranges *0* and *1* has range *01* as the intermediary state. Here we also expand this particular MuSSE-type model to allow for up to five hidden states and can be used to test hidden state character-dependent or character-independent diversification models, depending on how the different diversification parameters are set up (Table 1). In general, we include this particular set of models as a way of acknowledging that we really never know the “true” history of the characters or areas we observe. Therefore, there should be some way for the data to speak to scenarios that may be outside our *a priori* expectations with respect to geographic state speciation and extinction. Again, our argument follows the same logic as with the usage of hidden state SSE models. If a cladogenetic model is not the most adequate for our empirical data set, it is best to allow for a non-trivial “null” model (i.e., the anagenetic model) than to force a set of inadequate cladogenetic models to fit such data set. We refer to this class of models hereafter as “Anagenetic”.

**Table 1.** List of 18 fitted models. Columns show model category, model number, description of the model, and number of free parameters. Area-independent models (AIDiv) have no relationship between diversification rates and geographic areas. ‘full model’ indicates that all parameters of the model are free whereas ‘null model’ indicates that diversification and dispersion parameters are constrained to be equal among areas in the same hidden state category. If not ‘full model’ or ‘null model’, the description column lists the free parameters of the model. Models indicated with ‘+extirpation’ separate rates of range reduction from the extinction of endemic lineages.

| Category | Model | Description | free parameters |
| --- | --- | --- | --- |
| C<br>L<br>A<br>D<br>O<br>G<br>E<br>N<br>E<br>T<br>I<br>C | 1 | AIDiv - original GeoSSE, free dispersal | 4 |
|  | 2 | original GeoSSE, full model | 7 |
|  | 3 | AIDiv - GeoHiSSE, 3 rate classes, null model | 9 |
|  | 4 | GeoHiSSE, 2 rate classes, full model | 15 |
|  | 5 | AIDiv - GeoHiSSE, 5 rate classes, null model | 13 |
|  | 6 | AIDiv - GeoHiSSE, 2 rate classes, free dispersal | 7 |
| C +<br>L E<br>A X<br>D T<br>O I<br>G R<br>E P<br>N A<br>E T<br>T I<br>I O<br>C N | 7 | AIDiv - GeoSSE+extirpation, free dispersal | 6 |
|  | 8 | GeoSSE+extirpation, full model | 9 |
|  | 9 | AIDiv - GeoHiSSE+extirpation, 3 rate classes, null model | 11 |
|  | 10 | GeoHiSSE+extirpation, 2 rate classes, full model | 19 |
|  | 11 | AIDiv - GeoHiSSE+extirpation, 5 rate classes, null model | 15 |
|  | 12 | AIDiv - GeoHiSSE+extirpation, 2 rate classes, free dispersal | 9 |
| A<br>N<br>A<br>G<br>E<br>N<br>E<br>T<br>I<br>C | 13 | AIDiv - anagenetic GeoSSE, free dispersal | 6 |
|  | 14 | anagenetic GeoSSE, full model | 10 |
|  | 15 | AIDiv - anagenetic GeoHiSSE, 3 rate classes, null model | 11 |
|  | 16 | anagenetic GeoHiSSE, 2 rate classes, full model | 21 |
|  | 17 | AIDiv - anagenetic GeoHiSSE, 5 rate classes, null model | 15 |
|  | 18 | AIDiv - anagenetic GeoHiSSE, 2 rate classes, free dispersal | 9 |

*Simulations* ‐‐ We performed extensive simulations to test the behavior of the hidden state geographic state speciation and extinction (GeoHiSSE), in addition to our other model expansions. We evaluated models of area-dependent and independent diversification under a series of scenarios, including unequal frequencies between observed ranges and absence of cladogenetic events. We have relegated many of the details and tests to the Supplemental Materials. Briefly, we generated 100 trees containing 500 taxa for each simulation scenario. Thus, our results are relevant to trees of similar size or larger and we strongly suggest users to perform power analyses when using smaller trees. All analyses were carried out using new functions provided in the R package *hisse* (Beaulieu and O’Meara 2016). Code to replicate the simulations are also available on the *Supplementary material.*

For the first simulation scenario (scenario A), we simulated data using a homogeneous rate GeoSSE model with the speciation rate for one of the endemic areas (area *1*) set to be two times faster than the other two possible areas (*1* and *01*). This represented the simplest case for which the original GeoSSE model is known to be adequate (Goldberg et al. 2011). It also allowed us to test whether models with multiple hidden rate classes exert undue weight when rate heterogeneity is not actually present in the data. We found that even when the model set included a broad array of complex models, most of the model weight across all replicates for scenario A goes to the generating model (model 2; Figure S8). Furthermore, even when net diversification rates between endemic ranges are averaged across all models our estimates were congruent with the true values (Figure 4A). When we expressed speciation and extinction rates in terms of turnover rates (i.e., *s_i_* + *x_i_*) and extinction fraction (i.e., *x_i_*/*s_i_*), the rate estimates for each node in the tree are also centered on the true parameter values, independent of tree height (Figure S5).

**Figure 4.**
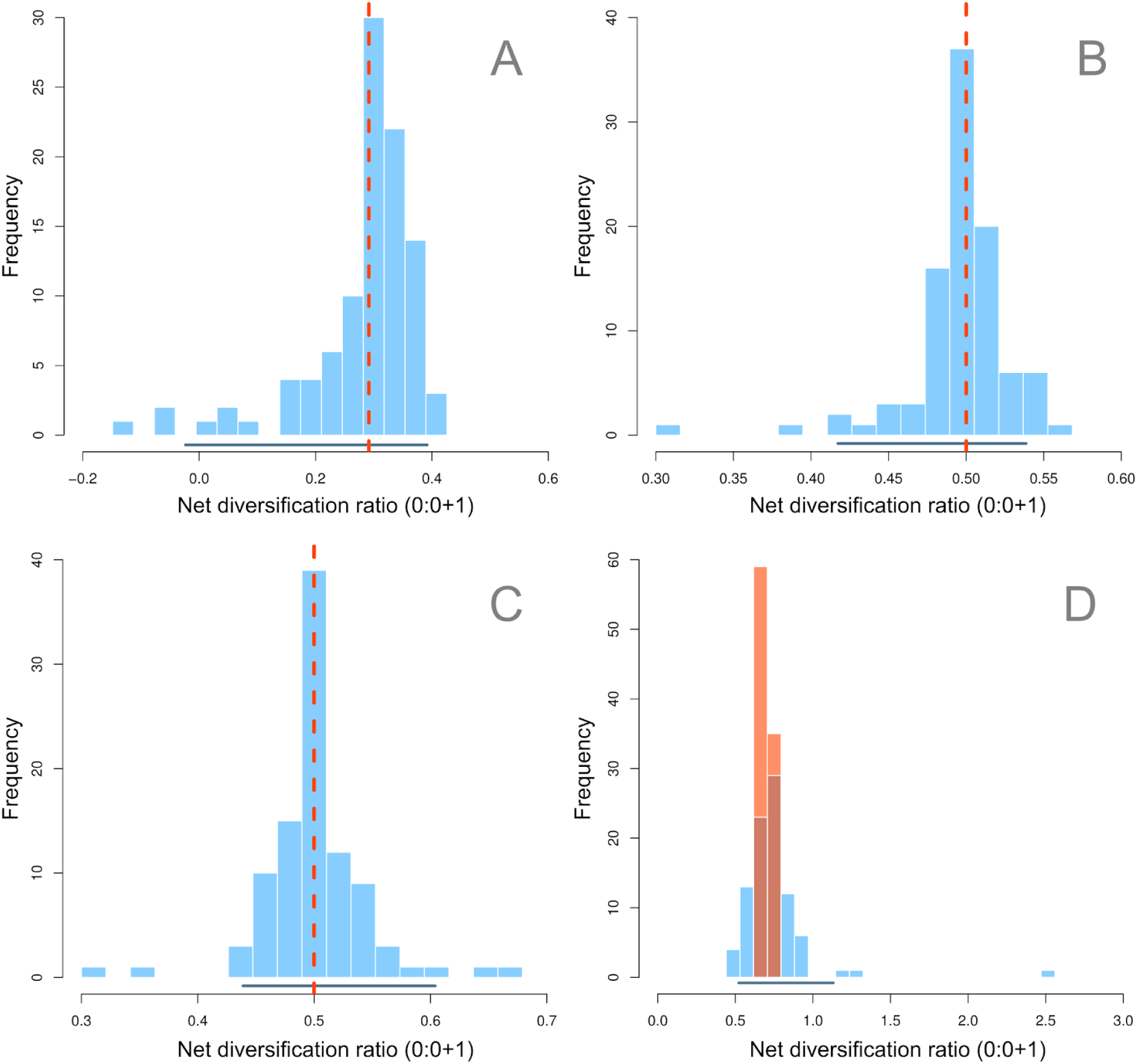
Effect of endemic ranges on net diversification estimated for simulation scenarios A to D. Plots show the distribution of ratios between net diversification rates for areas *0* and *0+1* computed for each simulation replicate. Red dashed vertical lines in plots A to C represent the true value for the ratios. Horizontal lines in the bottom show the 95% density interval for each distribution of parameter estimates. Plot D shows the distribution of true ratios in orange. Estimates are the result of model averaging across 18 different models using Akaike weights. See Table 1 for the list of models and Figure S3 for AIC weights.

In the second and third scenarios (scenarios B and C), we introduced three and five range-independent diversification regimes, respectively. The transition between diversification rate classes followed a meristic Markov model to emulate gradual changes in diversification rates (Figure S1). In both cases, we did not detect any meaningful differences in the net diversification rates between endemic areas (Figures 4B-C and S3), and the parameter estimates computed for each node and tip of the phylogeny are centered on the true values. Thus, our results show that area-independent GeoHiSSE models can accommodate the rate heterogeneity in the simulated trees without evoking an area-dependent diversification process when none was present. We do note, however, that even when we reduced the model set and fit only simple homogeneous GeoSSE models to the same data, there is an interesting effect in that the parameter estimates averaged across all models would still lead to an area-independent diversification interpretation (see *Appendix 2*).

Simulation scenario D represented an instance of the area-dependent model (ADDiv) model in which geography has an important effect on diversification across the phylogeny, but diversification rates vary within each range as a function of some unobserved “hidden” trait. The frequency of each hidden rate class stochastically varies across simulation replicates, and, therefore, there is no single true value for diversification rates. Nevertheless, results showed that GeoHiSSE was able to recover the correct direction in the relationship between net diversification rates of endemic areas in the presence of heterogeneity (Figures 4D, S5 and S6D).

We also studied two extreme cases with the objective of identifying odd behaviors when simulating data sets where 1) widespread ranges are rare or absent in extant species, and where 2) the evolution of areas are not tied to cladogenetic events. Under the original GeoSSE model, lineages need to pass through the widespread state before transitioning between endemic areas. If extant widespread lineages are rare or absent, the information to infer cladogenetic and dispersion events can become limited. To study this effect we first simulated datasets with widespread lineages as being rarely observed at the tips (see scenario E in Table S1). Results showed that low frequency of widespread lineages does not prevent our set of models from reaching meaningful estimates using model-averaging (Figure 5E). Alternatively, we simulated the case of jump dispersal events (i.e., direct transitions between endemic distributions). For this we used a GeoSSE model to simulate the data, but we allowed lineages to disperse between endemic areas without becoming widespread first (scenario F). [Note that none of the 18 models we used to estimate parameters throughout this study allow for any instance of jump dispersal events.] Our results showed no evidence for a meaningful bias in parameter estimates for both area-dependent diversification rates or speciation rates associated with cladogenetic events on widespread lineages (Figure 5F). In summary, our approach of model-averaging across a large set of candidate models does not appear sensitive to rare extant widespread areas.

**Figure 5.**
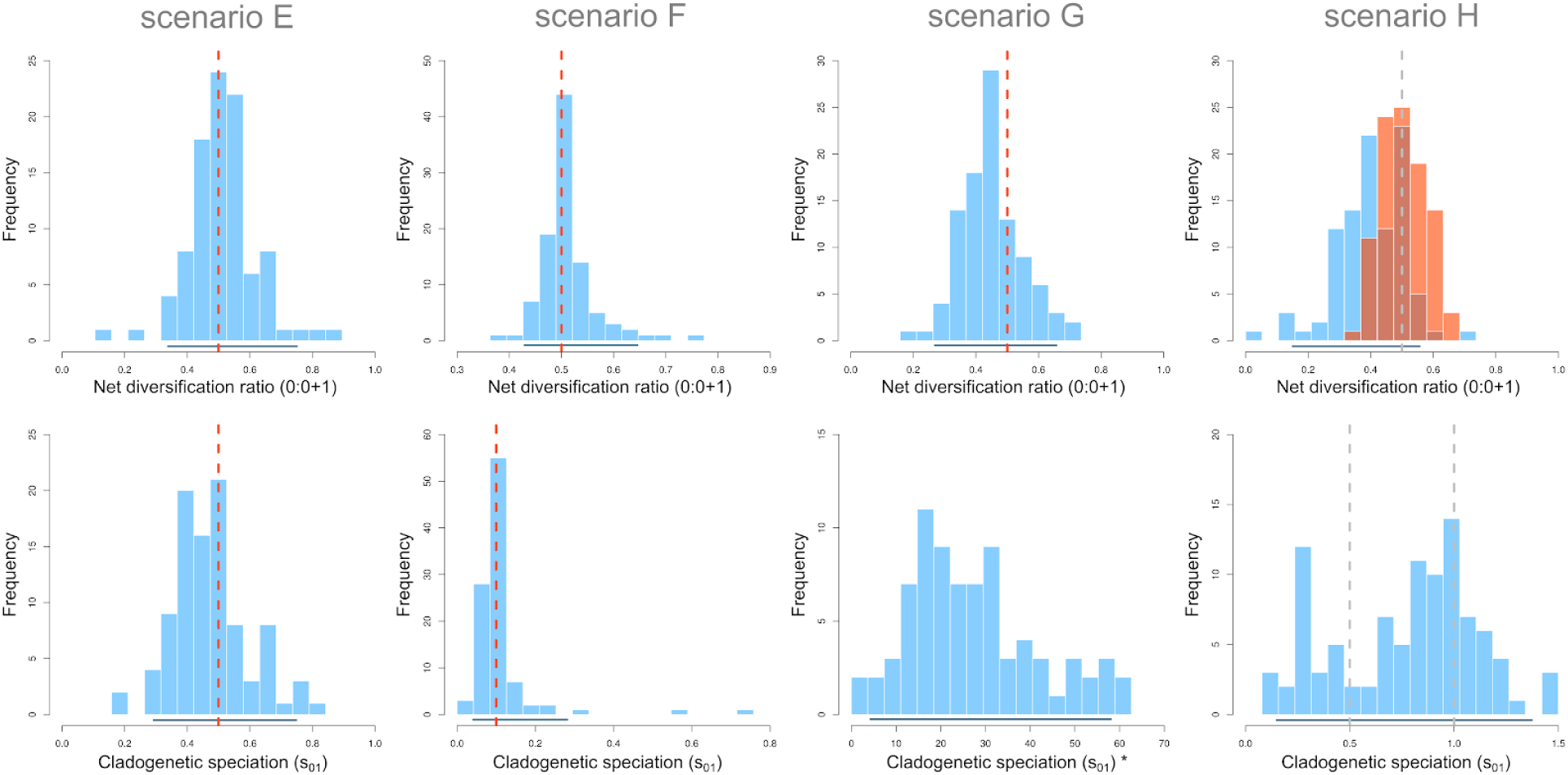
Net diversification ratios between endemic areas and cladogenetic speciation rates estimated for simulation scenarios E to F. Upper row shows the distribution of ratios between net diversification rates for areas *0* and *0+1* computed for each simulation replicate. Lower row shows the distribution of speciation rates associated with the widespread area (parameter *s_01_*) averaged across all tips (for E, F and H) or nodes (for G) of the phylogeny for each simulation replicate. Red dashed vertical lines represent the true value for the parameter. Grey dashed lines mark important reference points but are not the expected value for the quantities. Plot upper H shows the distribution of true ratios in orange (see main text). Horizontal lines in the bottom show the 95% density interval for each distribution of parameter estimates. The ‘∗’ marks results based on the average across nodes instead of tips (no data available at the tips). Estimates are the result of model averaging across 18 different models using Akaike weights. See Table 1 for the list of models and Figure S4 for AIC weights.

Finally, we explored the extreme possibility that the widespread range was never a part of the history of the clade (scenario G). When fit to our model set, the absence of widespread areas among the extant species produces estimates of the rates of cladogenetic speciation (*s_AB_*) that are highly uncertain (Figure 5G). These estimates are orders of magnitude higher than the rates of speciation associated with endemic regions. In contrast, estimates for the relative difference in net diversification rates between endemic areas did not show a strong bias (Figure 5G). This suggests that poor estimates of cladogenetic speciation would not strongly bias our conclusions about range-dependent diversification rates.

All previous scenarios assumed that cladogenetic events were important in the evolutionary history of the lineages. In order to consider the performance of the model when this is not the case, we generated datasets with transitions between areas restricted only to anagenetic dispersal events along the branches of the tree. The estimated difference in rates of diversification between endemic areas is larger than observed in any other simulation scenario (Figure 5H). Moreover, the absence of cladogenetic events makes estimates of cladogenetic speciation (*s_AB_*) uncertain, although raw parameter values are within the same order of magnitude of the true rates of diversification across the tree (grey lines in Figure 5H).

On the whole, our extensive simulation study shows that parameter estimates averaged across 18 models of area-independent and area-dependent evolution are robust to a wide variety of macroevolutionary scenarios likely to be observed in empirical datasets. We also show that important violations of the expected type of data modelled with GeoSSE, such as absence of widespread lineages or cladogenetic speciation events, are not enough to significantly hinder our interpretation of the evolutionary history of the group, even when there is large ambiguity in the estimate for some parameters of the model.

### Empirical example: Hemisphere-scale differences in conifer diversification

A further question is the performance of the GeoHiSSE model, as well as our extensions to the original GeoSSE model, in an empirical setting, where we do not know the generating model, and the tree and area designations could contain unforeseen errors or problematic and/or conflicting signals in the data. For these purposes, we focus our analyses on the evolutionary dynamics of movements between Northern and Southern Hemisphere conifers. There is evidence that the turnover rate — defined here as speciation + extinction rate — is generally higher for clades found exclusively in the Northern Hemisphere compared to clades found almost exclusively in the Southern Hemisphere (Leslie et al. 2012). The falling global temperatures throughout the Cenozoic, and concomitant movements of several major landmasses northwards, facilitated the emergence of colder, drier, and strongly seasonal environments within Northern Hemisphere regions (e.g., Zanazzi et al. 2007, Eldrett et al. 2009). This may have led to widespread extinction of taxa unable to survive in such environments and expansion of taxa able to thrive there (perhaps through isolated populations surviving to become new species rather than go extinct due to competition). The net effect across the clade would be an increase in speciation and extinction rates. Furthermore, the repeated cycles of range expansion and contraction due to glacial cycles would also promote isolation of populations leading both to speciation (due to allopatry), and possibly rapid extinction (due to small population size). The Southern Hemisphere, on the other hand, has maintained milder environments scattered throughout the region (e.g., Wilford and Brown 1994, McLoughlin 2001). It is important to note that these conclusions were supported by comparisons of branch length distributions and diversification models applied to various clades independently, which indicated heterogeneity among the taxonomic groups tested (see Leslie et al. 2012).

Using the expanded GeoSSE framework, we re-examined the hemisphere diversification differences within conifers proposed by Leslie et al. (2012). We combined geographic locality information from GBIF with an updated version of the dated conifer tree from Leslie et al. (2018) that improves taxon sampling relative to the analysis of Leslie et al. (2012). This new phylogeny contains 578 species, representing around 90% of the recognized extant diversity. We considered any locality having a maximum and minimum latitude >0 degrees as being Northern Hemisphere, <0 as Southern Hemisphere, and species with a maximum latitude >0 and a minimum latitude <0 were considered “widespread”. Such strict thresholds in latitude used to define ranges provide the ideal scenario in which hidden states may play an important role in understanding the diversification dynamics across the clades. Finally, we pruned the Pinaceae from our analysis and focus only on movements within the Cupressophyta, which includes the Cupressales (i.e., cypresses, junipers, yews, and relatives) and the Araucales (i.e., *Araucaria*, *Agatha*, podocarps, and relatives). The decision to remove Pinaceae from our analysis was based on the uncertain relationship of Gnetales to conifers. There remains the possibility that Gnetales is sister to conifers as a whole (e.g., the “Gnetifer” hypothesis; Chaw 1997, Burleigh and Mathews 2007), though most recent sequence analyses support Gnetales as sister to Pinaceae (e.g., the “Gnepine” hypothesis; Mathews 2006, Mathews 2009, Zhong et al. 2010). For these reasons, we focus our analyses on the Cupressophyta to ensure that our analyses reflect geographic range evolution within a monophyletic group.

Our final data set consisted of 325 species, with 146 species designated as Northern Hemisphere, 143 designated as Southern Hemisphere, and the remaining 36 species currently persisting in both areas. Our model set included the 18 models described in Table 1 and used in our simulations, as well as an additional 17 models. Briefly, we included AIDiv models that ranged from 2-5 hidden states rather than just those that equal the number of parameters in the ADDiv models (i.e., AIDiv models consisting of 3 and 5 hidden states), various MuSSE-type models that allowed and disallowed range contraction to be separate from lineage extinction, and a particular set of MuSSE-type models that disallowed speciation and extinction (i.e., the rates were set to zero) in the widespread regions to better mimic *anagenetic*-*only* range evolution. The entire set of models tested and their number of free parameters are described in Table S2.

The turnover rate differences, even after accounting for hidden states and the possibility of heterogeneity in area-independent diversification, supported the original findings of Leslie et al. (2012). That is, Northern Hemisphere species across Cupressophytes exhibited higher turnover rates relative to species occurring in the Southern Hemisphere (Figure 6). The majority of the model weight is comprised of estimates from three models (models 4, 10, and 32 described in Table 1 and Table S2), all of which assume a hidden state and character-dependent diversification, but differ in whether cladogenetic events have occurred or whether range contraction is separate from lineage extinction. The inference of a hidden state also implied that differences in the turnover rate were also clade-specific. Indeed, differences in the turnover rates within the Northern Hemisphere Cupressales (i.e., Taxaceae+Cupressaceae) have a turnover rate that is nearly 3 times higher than Southern Hemisphere species. Within the Araucariales (i.e., Araucariaceae+Podocarpaceae) the Northern Hemisphere rate is only 1.5 times higher than the species occurring in the Southern Hemisphere. Similarly, turnover rate is meaningfully higher in Southern Hemisphere Araucariales (0.152 events Myr^−1^) relative to the turnover rates among Southern Hemisphere members of the Cupressales (0.087 events Myr^−1^). Turnover rates are also higher among Northern Hemisphere species of Cupressales than Northern Hemisphere species of Araucariales, although these differences do not appear to be very meaningful (0.22 vs. 0.20 events Myr^−1^).

**Figure 6.**
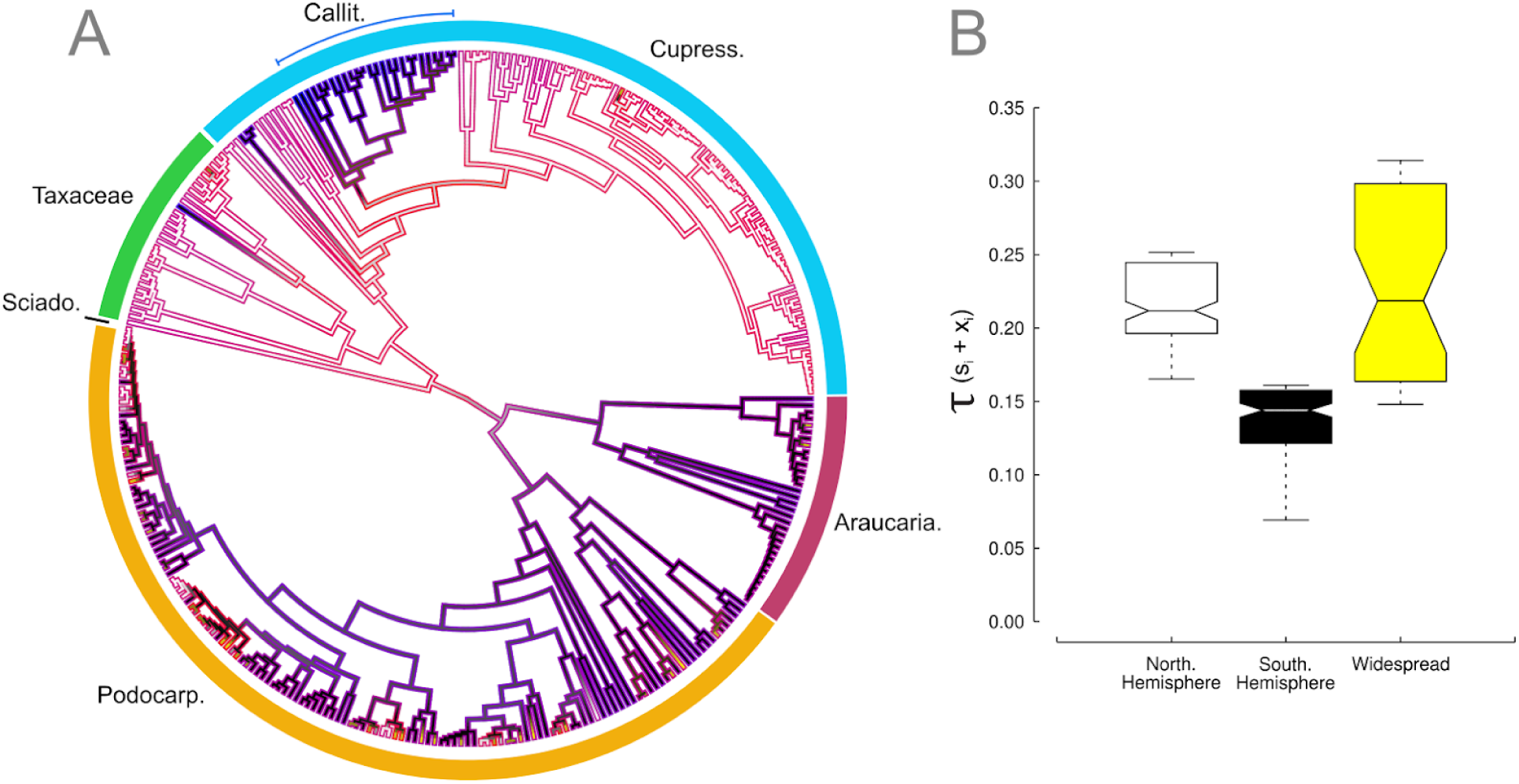
(A) Geographic area reconstruction of areas and turnover rates (i.e., τ = *s_i_* + *x_i_*) across a large set of models of Northern Hemisphere (white branches), Southern Hemisphere (black branches), and lineages in both (yellow branches) across Cupressophyta. The major clades are labeled and estimates of the most likely state and rate are based on the model-averaged marginal reconstructions inferred across a large set of models (see main text). The color gradient on a given branch ranges from the slowest turnover rates (blue) to the highest rates (red). (B) The distribution of turnover rates estimated for contemporary species currently inhabiting each of the geographic areas indicate that both Northern Hemisphere and widespread species generally experience higher turnover rates (i.e., more speciation and more extinction) relative to Southern Hemisphere species. Araucaria. = Araucariaceae, Callit. = Callitroids, Cupress. = Cupressaceae, Podocarp. = Podocarpaceae, Sciado. = Sciadopityaceae.

In cases where diversification rates (or any other rate of interest) are always higher in one particular observed state than the other for any hidden state, interpretation of the results is fairly straightforward (e.g., Figure 6). Mapping the model-averaged diversification rates to the phylogenetic tree can also provide important, and often insightful, phylogenetic context to among species variation in parameter estimates. However, in some instances, ignoring rate differences between combinations of observed and hidden traits could be problematic. Whether or not one observed state leads to higher diversification rates than the other could depend on the magnitude of the rate differences and how much time is spent in each hidden state ‐‐ that is, when the rate of diversification for *0A*> 1*A*> *1B* > *0B* the average mean rate for *0* could be equal, higher than, or lower than the rate for state *1.*

As a solution to this, we recommend model-averaging the “effective trait ratio”, which is the expected proportion of each observed state given the estimated parameters, tree depth, and the root weights under each of the models in the set (see *Appendix 1*). This provides an intuitive complement to examining just rate differences among observed states. In other words, even in cases where *0A*> *1A*> *1B* > *0B*, and where the diversification rate of state *0* is more or less the same as state *1*, we could still find that, say, 75% of species are expected to be in state *0* due to the interaction with the hidden character. Note that this may differ radically from the empirical frequency, which is based on one realization of the process. Perhaps an unlikely series of changes early in the tree led to more taxa in state *1* than would be expected. Likewise, under a standard BiSSE analysis, it is possible to have more taxa in state *1* even if the net diversification rate in state *1* and transition rate to state *1* are lower than the equivalent rates for *0.* Given this scenario, one should conclude that there is signal for a trait-dependent process with observed state *0* having a positive effect on diversification, despite the lack of a consistent direction in the diference between hidden classes of observed traits. In the case of Cupressophytes, the model-averaged ratios indicated that roughly 52.3% (±9.6%) of species are expected to occur in the Southern Hemisphere, with 35.4% (± 12.9%) of conifers are expected to be in the Northern Hemisphere, and 12.3% (±4.4%) of the species widespread across both regions. These expectations are largely congruent with the estimates of turnover rates between Northern and Southern Hemisphere, where the increased “boom and bust” dynamics in the Northern Hemisphere produces diversity within the region at much lower levels than in the Southern Hemisphere.

## Caveats

The GeoHiSSE family of models are reasonable approaches for understanding diversification when there can be many processes at play. However, it is important that we emphasize that, like all models, they are far from perfect. First, the hidden traits evolve under a continuous time Markov process, which is reasonable for heritable traits that affect diversification processes. Of course, not all heterogeneity arises in such a way. For instance, when an asteroid impact throws up a dust cloud, or causes a catastrophic fire, every lineage alive at that time is affected simultaneously. Their ability to survive may come from heritable factors, but the sudden shift in diversification caused by an exogenous event like this appears suddenly across the tree, in a manner not yet incorporated in these models. Similarly, a secular trend affecting all species is not part of this model, but is in others (Morlon et al. 2011), and they also do not incorporate factors like a species “memory” of time since last speciation (Alexander et al. 2015), or even a global carrying capacity for a clade that affects the diversification rate (Rabosky and Glor 2010). These models require large numbers of taxa to be effective. Investigating the evolution of compelling but small clades like the great apes, baleen whales, or the Hawaiian silverswords can be attempted with these approaches, but results are likely to be met with fairly uncertain parameter estimates. It it possible to run these approaches over a cloud of trees. This will not compensate for biases or errors in the trees, such as if trees were inferred using a Yule prior and lack enough information to overwhelm the prior, if sequencing errors lead to overly long terminal branches, or if reticulate evolution is present and not taken into account. Most of these caveats are not limited to HMM models of diversification, but this reminder may serve to reduce overconfidence in results.

## Conclusions

In practice, SSE models are generally only considered in a hypothesis rejection framework, namely, reject a null model where there is no trait-dependent diversification and thus accept an alternate model where rates depend on traits. The term “reject” can mean formal rejection using a likelihood ratio test, but it often takes the form of selecting the alternate model under AIC (despite warnings from Burnham and Anderson 2002). Rabosky and Goldberg (2015) vividly showed problems with BiSSE when interpreted this way. While accurately critiquing how scientists use BiSSE models, their results point more to deficiencies in how we biologists use statistics rather than a particular problem with the SSE models *per se* (see Beaulieu and O’Meara 2016).

As scientists, we have all learned to recognize and worry about Type I errors, and so it is not surprising that this has remained the *status quo* when examining model behavior. In many cases reviewers may insist upon testing for the “best” model as a confirmatory approach, even when, as we show here, model-averaging has good performance and is robust to deviations from model assumptions (see similar comment on reviewer insistence in Ree and Sanmartín 2018). But, if we are to proceed down this road with SSE models, a model where the trait *and* the tree both evolved under constant rates of speciation and extinction and, sometimes, even under constant state transition rates is *not* the “null” model. Shifts in rates of diversification are ubiquitous across the tree of life for many reasons (mass extinctions, adaptive radiations, changing biogeography, available niches, etc.) and it is a safe bet to assume nearly any empirical tree will not show perfectly constant speciation and/or extinction rates. In the case of an empirical tree with trait-independent diversification, the null model (of constant rates for everything) and the alternative model (that traits affect diversification) are both wrong. So, which model should a good test choose?

In our view, even when presented with a equally complex alternative, the focus on model rejection still remains problematic. In certain areas of biology, we seem to stop after rejecting a null model that we already knew was false, though to be fair, of course, it is useful to get information about whether or not an effect might be present. But scientists should go beyond this to actually look at parameter estimates. Suppose we find the diversification rate of red flowers are higher than yellow flowers. Is it 1% higher, or is it 300% higher? The answer could have biologically very different implications, to the extent that rejecting or not the null model becomes largely irrelevant. Such concerns are much more common in other applications, particularly with linear regression models. However, the same care of checking how well the regression line passes through the data points and interpreting *R*^2^, among other tests, have counterparts in phylogenetic comparative models and are equally relevant.

Here we show that hidden state modelling (HMM) is a general framework that has the potential to greatly improve the adequacy of any class of SSE models. The inclusion of hidden states are a means for testing hypotheses about unobserved factors influencing diversification in addition to the observed traits of interest. We emphasize, however, that they should not be treated as a separate class of SSE models, but instead viewed as complementary and should be included as part of a set of models under evaluation. They also represent a straightforward approach to incorporating different types of unobserved heterogeneity in phylogenetic trees than a simple single rate category model is able to explain. For example, our expanded suite of GeoSSE models allows accommodating heterogeneity in the diversification process as it relates to geographic areas. The GeoHiSSE models introduced here can be applied to study rates of dispersion and cladogenesis as well as to perform ancestral area reconstructions, thus being a suitable alternative to avoid the shortcomings of DEC and DEC+J models (Ree and Sanmartín 2018). Moreover, the area-independent diversification models (AIDiv) appear adequate to explain shifts in diversification regimes unrelated to geographic ranges, and demonstrate that the area-dependent models (ADDiv) can successfully estimate the impact of geographic areas on diversification dynamics when such a signal is present in the data.

Many phylogenetic comparative methods have been under detailed scrutiny recently (e.g., Maddison and FitzJohn 2015, Rabosky and Goldberg 2015, Cooper et al. 2016a, Cooper et al. 2016b, Adams and Collyer 2017, Ree and Sanmartín 2018). This is certainly a worthy endeavor given that all models will fail under certain conditions, and some models have innate flaws that render them unwise to use. One response (e.g., Rabosky and Huang 2016, Rabosky and Goldberg 2017, Adams and Collyer 2017, Harvey and Rabosky 2018) has been to move to “semi-” or “non-parametric” approaches, some of which do incorporate models internally, but with an end result that is a non-parametric test. The issue with this is that they move entirely away from estimating parameters and the entire exercise becomes rejecting null models that we never really believed in. Such methods, in our view, provide very limited insights as they only show that the null model is wrong, with appropriate Type I error, which is not the same as showing the alternate is a correct model of the world. A more fruitful approach may be improving upon the existing models and better communicating when methods can and cannot be applied (Cooper et al. 2016a). We should stop trying to prove that our data cannot be explained by naive, biologically-blind null models none of us believe and, instead, fit appropriate models such that we can learn about meaningful patterns and processes across the tree of life.

## Acknowledgements

We thank members of the Beaulieu, O’Meara, and Alverson labs for their comments and for general discussions of the ideas presented here; we especially thank Teo Nakov and James Boyko. We would also like to specifically thank Andrew Alverson and Stacey Smith for their insightful critiques and helpful edits on an earlier version of this manuscript. DSC would also like to thank the Coordenação de Aperfeiçoamento de Pessoal de Nível Superior (CAPES: 1093/12-6) for the opportunity to work on this project.

## Appendix 1

*Effective trait ratios under GeoHiSSE* ‐‐ We can use parameters estimated under a given model to determine the expected frequency of each observed area and hidden state combination across a long stretch of evolutionary time. These equilibrium frequencies are often used in a variety of ways, most notably as weights in the likelihood calculation at the root (see Goldberg et al. 2011). We rely on them to compliment the examination of rate differences among the observed ranges. In the main text we describe a situation in which the rate of observed state *0* is more or less the same as state *1*, but given the interaction with the hidden characters in the model we may find that 75% of species are expected to be in area *0* over some specified length of evolutionary time.

We follow Maddison et al. (2007) and track the number of lineages in area 0, *n*_0_, the number of lineages in area *1*, *n*_1_ and the number of lineages in the widespread area *01*, *n*_01_, over some length of evolutionary time, *T.* The index, *i*, denotes the possible hidden states (*A*, *B*, …, *i*) that each observed state is associated. Given all the possible events that could occur across any given interval of time, we obtain the expected number of species for each area through the following ordinary differential equations:

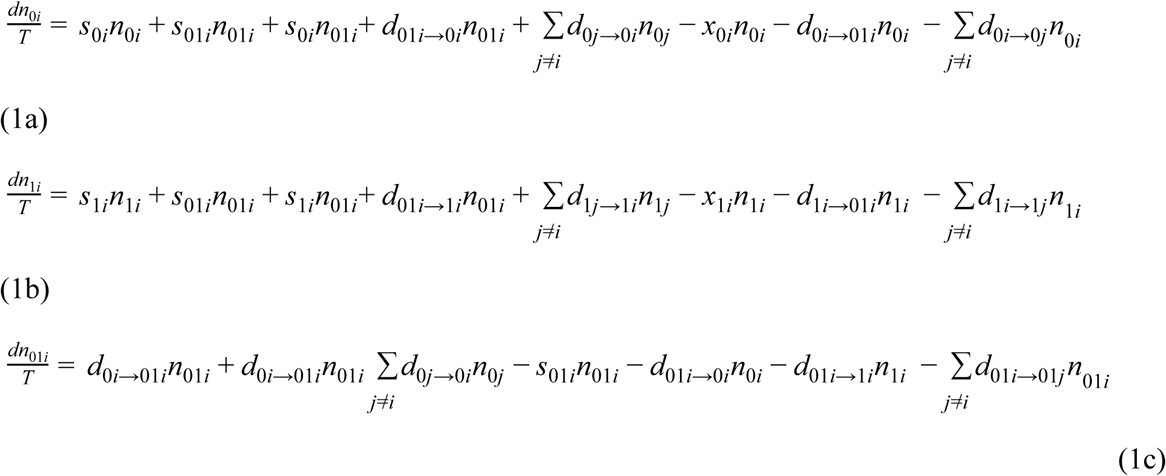

Note that we use *d*_0l*i*→0*i*_ and *d*_01*i*→1*i*_ to denote instances in which range contraction is separate from lineage extinction, *x*_01*i*→0*i*_. and *x*_1_. However, if the model assumes range contraction is governed by the same parameter value as lineage extinction, then we simply set *d*_01*i*→1*i*_ = *x*_1_ and *d*_01*i*→1*i*_ = *x*_0_. We also note that when there are no hidden states these equations reduce exactly to the equilibrium frequencies under the original GeoSSE formulation. The initial conditions are set according to the state at the root. In our case, as a means of accounting for the uncertainty in the starting state we rely on the likelihood that each area gave rise to the data (FitzJohn et al. 2009). Once the number of lineages are determined after a specified *T* we then sum the frequencies for each observed area across each hidden state and normalize them so that the observed area frequencies sum to 1.

The above equations assume explicitly that the birth-death process directly impacts range evolution through cladogenetic events, which are not allowed if the underlying model is a MuSSE-type model. Thus, in the MuSSE-type case, we rely on the following ordinary differential equations:

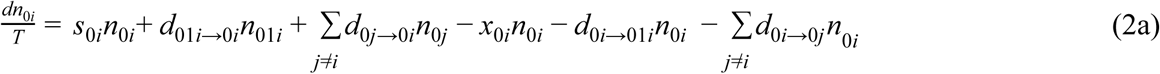

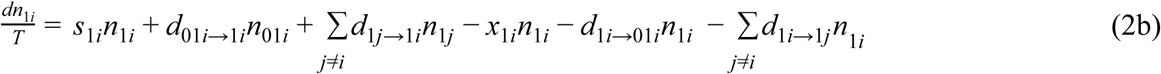

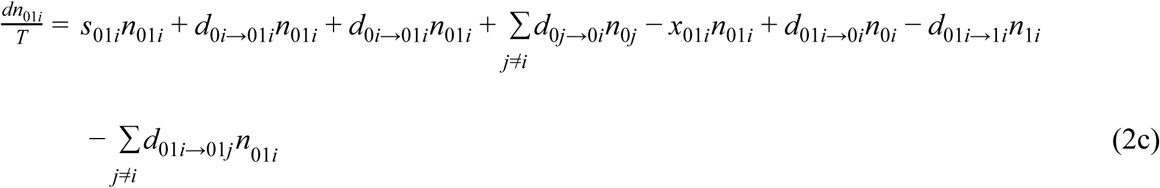

Author contributions
JMB and BCO conceived the hidden-Markov approach applied to SSE models. DSC, BCO, and JMB designed simulations, derived models, implemented model averaging and R code. DSC conducted simulations. JMB conducted empirical analyses. DSC, BCO, and JMB wrote the manuscript.

## References

Adams, D.C., and M.L. Collyer. 2017. Multivariate phylogenetic comparative methods: evaluations, comparisons, and recommendations. Syst. Biol. 67:14–31.

Alexander, H.K., A. Lambert, and T. Stadler. 2015. Quantifying age-dependent extinction from species phylogenies. Syst. Biol. 65:35–50.

Alfaro, M.E., F. Santini, C. Brock, H. Alamillo, A. Dornburg, D. L. Rabosky, G. Carnevale, and L.J. Harmon. 2009. Nine exceptional radiations plus high turnover explain species diversity in jawed vertebrates. Proc. Natl. Ac. Sci., USA 106:13410–13414.

Beaulieu, J.M., and B.C. O’Meara. 2016. Detecting hidden diversification shifts in models of trait-dependent speciation and extinction. Syst. Biol. 65:583–601.

Berkson, J. 1938. Some difficulties of interpretation encountered in the application of the chi-square test. J. Am. Stat. Assoc. 33:526–536.

Burleigh, J.G. and S. Mathews. 2007. Assessing systematic error in the inference of seed plant phylogeny. Int. J. Plant Sci. 168:125–135.

Burnham, K.P., Anderson D.R. 2002. Model selection and multimodel inference: a practical information-theoretic approach. New York:Springer.

Butler, M.A., and A.A. King. 2004. Phylogenetic comparative analysis: a modeling approach for adaptive evolution. Am. Nat. 164:683–695.

Chaw, S.M., Zharkikh A., Sung H.M., Lau T.C., Li W.H. 1997. Molecular phylogeny of extant gymnosperms and seed plant evolution: analysis of nuclear 18S rRNA sequences. Mol. Biol. Evol. 14:56–68.

Cooper, N., G.H. Thomas, and R.G. FitzJohn. 2016a. Shedding light on the ‘dark side’ of phylogenetic comparative methods. Methods Ecol. Evol. 7:693–699.

Cooper, N., G.H. Thomas, C. Venditti, A. Meade, and R.P. Freckleton. 2016b. A cautionary note on the use of Ornstein Uhlenbeck models in macroevolutionary studies. Biol. J. Linn. Soc. 118:64–77.

Dalling, J.W., P. Barkan, P.J. Bellingham, J.R. Healey, and E.V.J. Tanner. 2011. Ecology and Distribution of Neotropical Podocarpaceae. L. Cernusak, B. Turner (Eds.), Ecology of the Podocarpaceae in Tropical Forests, Smithsonian Institution, Washington (2011), pp. 43–56.

Eastman, J.M., M.E. Alfaro, P. Joyce, A.L. Hipp, and L.J. Harmon. 2011. A novel comparative method for identifying shifts in the rate of character evolution on trees. Evolution 65:3578–3589.

Eldrett, J.S., D.R. Greenwood, I.C. Harding, and M. Huber. 2009. Increased seasonality through the Eocene to Oligocene transition in northern high latitudes. Nature 459:969–973.

FitzJohn, R.G. 2012. Diversitree: comparative phylogenetic analyses of diversification in R. Methods Ecol. Evol. 3:1084–1092.

FitzJohn, R.G., W.P. Maddison, and S.P. Otto. 2009. Estimating trait-dependent speciation and extinction rates from incompletely resolved phylogenies. Syst. Biol. 58:595–611.

FitzJohn, R.G. 2010. Quantitative traits and diversification. Syst. Biol. 59:619–633.

Goldberg, E.E., and B. Igić. 2012. Tempo and mode in plant breeding system evolution. Evolution, 66:3701–3709.

Goldberg, E.E., L.T. Lancaster, and R.H. Ree. 2011. Phylogenetic inference of reciprocal effects between geographic range evolution and diversification. Syst. Biol. 60:451–465.

Green, P. J. 1995. Reversible jump Markov chain monte carlo computation and Bayesian model determination. Biometrika, 82:711–732.

Harvey MG, Rabosky DL. 2018. Continuous traits and speciation rates: Alternatives to state-dependent diversification models. Methods Ecol. Evol. doi:10.1111/2041-210X.12949

Huang, D., E.E. Goldberg, L.M. Chou, and K. Roy. 2018. The origin and evolution of coral species richness in a marine biodiversity hotspot. Evolution, 72:288–302.

Kirk, R.E. 1996. Practical significance: A concept whose time has come. Educ. Psychol. Meas. 56:746–759.

Kunzmann, L. 2007. Araucariaceae (Pinopsida): Aspects in palaeobiogeography and palaeobiodiversity in the Mesozoic. Zool. Anz. 246:257–277.

Leslie, A.B., J.M. Beaulieu, H.S. Rai, P.R. Crane, M.J. Donoghue, and S. Mathews. 2012. Hemisphere-scale differences in conifer evolutionary dynamics. Proc. Natl. Ac. Sci., USA. 109:16217–16221.

Leslie, A.B., J.M. Beaulieu, G. Holman, C.S. Campbell, W. Mei, L.R. Raubeson, and S. Mathews. 2018. An overview of extant conifer phylogeny from the perspective of the fossil record. Am. J. Bot. In press.

Maddison, W.P., and R.G. FitzJohn. 2015. The unsolved challenge to phylogenetic correlation tests for categorical characters. Syst. Biol. 64:127–136.

Maddison, W.P., P.E. Midford, and S.P. Otto. 2007. Estimating a binary character’s effect on speciation and extinction. Syst. Biol. 56:701–710.

Magnuson-Ford, K., and S.P. Otto. 2012. Linking the investigations of character evolution and species diversification. Am. Nat. 180:225–245.

Mathews, S. 2006. Phytochrome-mediated development in land plants: red light sensing evolves to meet the challenges of changing light environments. Mol. Ecol. 15:3483–3503.

Mathews, S. 2009. Phylogenetic relationships among seed plants: persistent questions and the limits of molecular data. Am. J. Bot. 96:228–236.

Matzke, N.J. 2014. Model selection in historical biogeography reveals that founder-event speciation is a crucial process in island clades. Syst. Biol. 63(6):951–970.

McLoughlin, S. 2001. The breakup history of Gondwana and its impact on pre-Cenozoic floristic provincialism. Aust. J. Bot. 49:271–300.

Morlon, H., T.L., Parsons, and J.B., Plotkin. 2011. Reconciling molecular phylogenies with the fossil record. Proc. Natl. Ac. Sci., USA. 08(39):16327–16332.

Nee, S., R.M., May, and P.H., Harvey. 1994. The reconstructed evolutionary process. Philos.Trans. Royal Soc. B. 344:305–311.

O’Meara, B.C., and J.M. Beaulieu. 2016. Past, future, and present of state-dependent models of diversification. Am. J. Bot. 103:1–4.

Pagel, M. 1994. Detecting correlated evolution on phylogenies: a general method for the comparative analysis of discrete characters. Proc. Royal Soc. B. 255:37–45.

Posada, D. 2008. jModelTest: phylogenetic model averaging. Mol. Biol. Evol. 25:1253–1256.

Rabosky, D. L. 2007. LASER: A Maximum Likelihood Toolkit for Detecting Temporal Shifts in Diversification Rates From Molecular Phylogenies. Evol. Bioinform. 2:247–250.

Rabosky, D.L. 2014. Automatic detection of key innovations, rate shifts, and diversity-dependence on Phylogenetic Trees. PLoS ONE, 9:e89543.

Rabosky, D.L., and R.E. Glor. 2010. Equilibrium speciation dynamics in a model adaptive radiation of island lizards. Proc. Natl. Ac. Sci., USA. 107:22178–22183.

Rabosky, D.L., and E.E. Goldberg. 2015. Model inadequacy and mistaken inferences of trait-dependent speciation. Syst. Biol. 64:340–355.

Rabosky, D.L., and E.E. Goldberg. 2017. FiSSE: A simple nonparametric test for the effects of a binary character on lineage diversification rates. Evolution, 71:1432–1442.

Rabosky, D.L., and H. Huang. 2016. A robust semi-parametric test for detecting Trait-Dependent diversification. Syst. Biol. 65:181–193.

Rabosky, D.L., and I.J. Lovette. 2008. Density dependent diversification in North American wood-warblers. Proc. Royal Soc. B. 275:2363–2371.

Ree, R.H., and I. Sanmartín. 2018. Conceptual and statistical problems with the DEC+J model of founder-event speciation and its comparison with DEC via model selection. J. Biogeogr. doi:10.1111/jbi.13173.

Ree, R.H., and S. Smith. 2008. Maximum likelihood inference of geographic range evolution by dispersal, local extinction, and cladogenesis. Syst. Biol. 57(1):4–14.

Rolland, J., F.L. Condamine, F. Jiguet, H. Morlon. 2014. Faster speciation and reduced extinction in the tropics contribute to the mammalian latitudinal diversity gradient. PLoS Biology, 12:e1001775.

Schluter, D., T. Price, A.Ø. Mooers, and D. Ludwig. 1997. Likelihood of Ancestor States in Adaptive Radiation. Evolution, 51:1699–1711.

Stadler, T. 2011. Mammalian phylogeny reveals recent diversification rate shifts. Proc. Natl. Ac.Sci., USA. 108:6187–6192.

Wilford, G.E. and P.J. Brown. 1994. History of the Australian Vegetation, ed. R.S. Hill, Cambridge University Press, Cambridge, pp. 5–13.

Yang, Z., S. Kumar, and M. Nei. 1995. A new method of inference of ancestral nucleotide and amino acid sequences. Genetics, 141:1641–1650.

Yang, Z., and R. Nielsen. 2008. Mutation-Selection Models of Codon Substitution and Their Use to Estimate Selective Strengths on Codon Usage. Mol. Biol. Evol. 25:568–579.

Zanazzi, A., M.J. Kohn, B.J. McFadden, and D.O. Terry, Jr. 2007. Large temperature drop across the Eocene-Oligocene transition in central North American. Nature, 445:639–642.

Zhong, B., O. Deusch, V.V. Goremykin, D. Penny, P.J. Biggs, R.A. Atherton, S.V. Nikiforova, and P.J. Lockhart. 2010. Systematic error in seed plant phylogenomics. Genome Biol. Evol. 3:1340–1346.

